# Imaging Synaptic Vesicle Protein SV2C with ^18^F-UCB-F: An In Vitro Autoradiography and In Vivo NHP PET Study

**DOI:** 10.64898/2026.03.08.710088

**Authors:** S. Nag, V. C. Sousa, R. Zou, A. F. Morén, P. Datta, Y. K. Meynaq, A. Valade, C. Vermeiren, P. Motte, J. Mercier, H. Ågren, C. Halldin, A. Varrone

## Abstract

The synaptic vesicle protein SV2C, predominantly found in the basal ganglia, has been associated with Parkinson’s disease through genetic studies. It plays a crucial role in regulating dopamine release and has been shown to be disrupted in PD animal models and brain tissues from PD patients. In the context of PD-related synaptopathy, SV2C may serve as a potential imaging target for monitoring disease progression and response to treatment.

[^18^F]UCB-F is a radioligand binding to SV2C developed by UCB. Preliminary autoradiography and PET studies in rats showed that [^18^F]UCB-F displays a brain distribution consistent with the expression of SV2C *in vitro* but does not display any specific binding *in vivo*. This study was therefore designed to further investigate the affinity and selectivity of [^18^F]UCB-F for SV2C and to examine the *in vitro* and *in vivo* properties of the radioligand in non-human primates. *In vitro* binding studies were performed to measure the affinity of UCB-F to SV2A, SV2B, and SV2C. Insilico modeling was used to assess the binding mode and energy of UCB-F. Autoradiography studies on rat and non-human primate (NHP) brain tissues were performed to confirm that [^18^F]UCB-F showed similar distribution in rat and NHP tissue. Finally, PET studied in NHPs were performed to examine the *in vivo* pharmacokinetic properties of [^18^F]UCB-F.

[^18^F]UCB-F was successfully synthesized from the corresponding precursor with high yield. Autoradiography on brain slices from rats and NHPs demonstrated specific binding of [^18^F]UCB-F in the pallidum, striatum, substantia nigra, and brainstem, consistent with the known brain expression of SV2C. In NHPs, [^18^F]UCB-F rapidly crossed the blood-brain barrier, reaching peak uptake values of 2.8 %ID in NHP1 and 2.1 %ID in NHP2 at 4 minutes post-injection. The tracer wasrapidly washed out from the brain, with no clear regional distribution. Radiometabolite analysis revealed the formation of only more polar radiometabolites, with approximately 15% of unchanged radioligand remaining in plasma at 15 minutes post-injection.

*In vitro* and *in-silico* studies demonstrated that the affinity of [^18^F]UCB-F decreased by approximately one factor of magnitude with increase of temperature from 4° to 37° C. This temperature-related decrease of the affinity for SV2C together with rapid *in vivo* radiometabolism might explain the discrepancy between *in vitro* and *in vivo* performance of [^18^F]UCB-F. Overall, these findings suggest that [^18^F]UCB-F is not a suitable PET radioligand for imaging SV2C. Further research is needed to identify alternative candidates with improved in vivo stability and brain retention.

## Introduction

Synaptic vesicle glycoprotein 2C (SV2C) is a synaptic vesicle membrane protein in the SV2 family involved in vesicle trafficking and vesicle-filled neurotransmitter release^1^. The three isoforms SV2A, SV2B, and SV2C show differences in the regional brain expression in the brain and diverse functions^2, 3^. In contrast to the broadly located SV2A, SV2C has a narrower, cell-type specific distribution with strong enrichment in the basal ganglia (caudate, putamen, globus pallidus, and substantia nigra) in dopaminergic cells and GABAergic projections. Preclinical and human tissue studies have shown that SV2C is involved in dopaminergic transmission and is affected in Parkinson’s disease (PD) ^4-6^. Studies on SV2C knock-out mice and animal models of PD show decreased dopamine release and alterations to presynaptic vesicle dynamics, with impaired motor behaviors due to deficits in dopaminergic signaling^7, 8^. Studies on brain tissue from PD donors showed disruption and reduced expression of SV2C^4^. Overall, these findings suggest that SV2C is an attractive molecular target for *in vivo* synaptic imaging with positron emission tomography (PET). We hypothesize that SV2C PET can be used as imaging tool for early detection of synaptic dysfunction and monitoring of synaptic changes in neurodegenerative disorders, particularly PD.

SV2A PET radioligands such as [^11^C]UCB-J^9, 10^ and [^18^F]SynVesT-1^11^, serve as imaging markers to study synaptic density, since SV2A is uniformly and highly expressed in the brain. SV2C has a lower density than SV2A and as restricted brain distribution. Therefore, the development of SV2C PET radioligands require compound optimization of affinity for SV2C and strong selectivity over SV2A/SV2B, as well as feasibility for labelling with ^11^C or ^18^F and suitable *in vivo* pharmacokinetic properties.

[^18^F]UCB-F is a PET radioligand for SV2C that has been developed by UCB and previously evaluated in rats using autoradiography and *in vivo* PET. On autoradiography [^18^F]UCB-F displayed a brain distribution consistent with the expression of SV2C. However, *in vivo* PET imaging showed no evidence of specific binding of [^18^F]UCB-F in the rat brain. The aim of this study was to further characterize [^18^F]UCB-F as potential SV2C PET radioligand. Specific objectives were: 1) To assess the in vitro affinity and selectivity to SV2C by in vitro binding studies and *in silico* modeling of binding mode and energy of UCB-F for SV2C; 2) To radiolabel [^18^F]UCB-F and to assess its *in vitro* binding in rat and non-human primate (NHP) brain tissue; 3) To assessthe *in vivo* pharmacokinetic properties of [^18^F]UCB-F in NHPs, with PET.

## Results and discussions

### In silico modelling

*In-silico* modelling was used to investigate the effect of temperature on UCB-F binding to SV2C. The initial structure of SV2C was generated using homology modelling, followed by structural relaxation through MD simulations. As shown in Figure 1, UCB-F adopts a consistent and welldefined binding conformation within the SV2C binding pocket, stabilized by two key hydrogen bonds. These conserved interactions contribute to anchoring the ligand and maintaining its orientation, suggesting that hydrogen bonding may play a critical role in supporting the stability of the UCB-F–SV2C complex.

**Figure 1.**
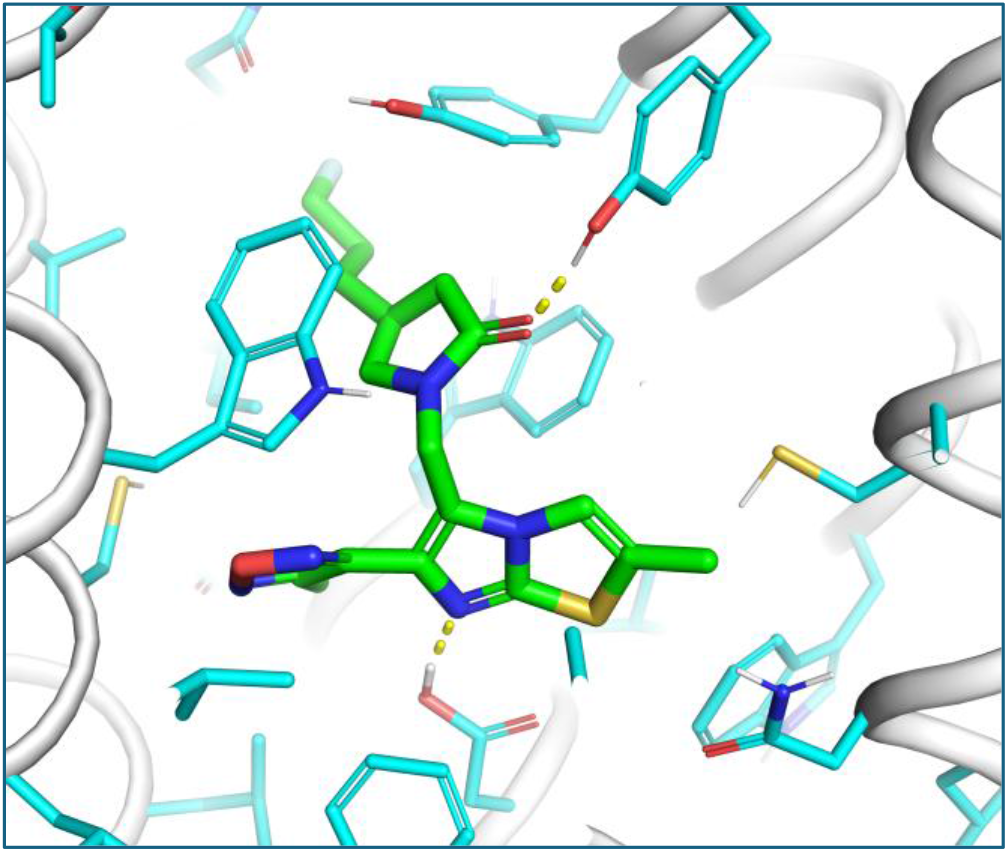
Molecular binding interactions within the SV2C–UCB-F complex. The protein backbone is shown as gray helices, and the ligand is depicted in stick representation with carbon atoms colored cyan and green, nitrogen in blue, and oxygen in red. Yellow dashed lines represent hydrogen bonds formed between the ligand and surrounding protein residues.

Molecular dynamics simulations were performed on the protein–ligand complexes at different temperatures to evaluate both the conformational stability and dynamic behaviour of the predicted binding poses. As shown in Figure 2, the temporal evolution of ligand RMSD was monitored throughout the 100 ns simulation trajectories. Across all replicates, RMSD values consistently remained below 2.5 Å, indicating that the initial binding conformations were stably maintained over time. These results demonstrate the robustness of the predicted binding mode and confirm that UCB-F remains well anchored within the binding pocket under the simulated conditions.

**Figure 2.**
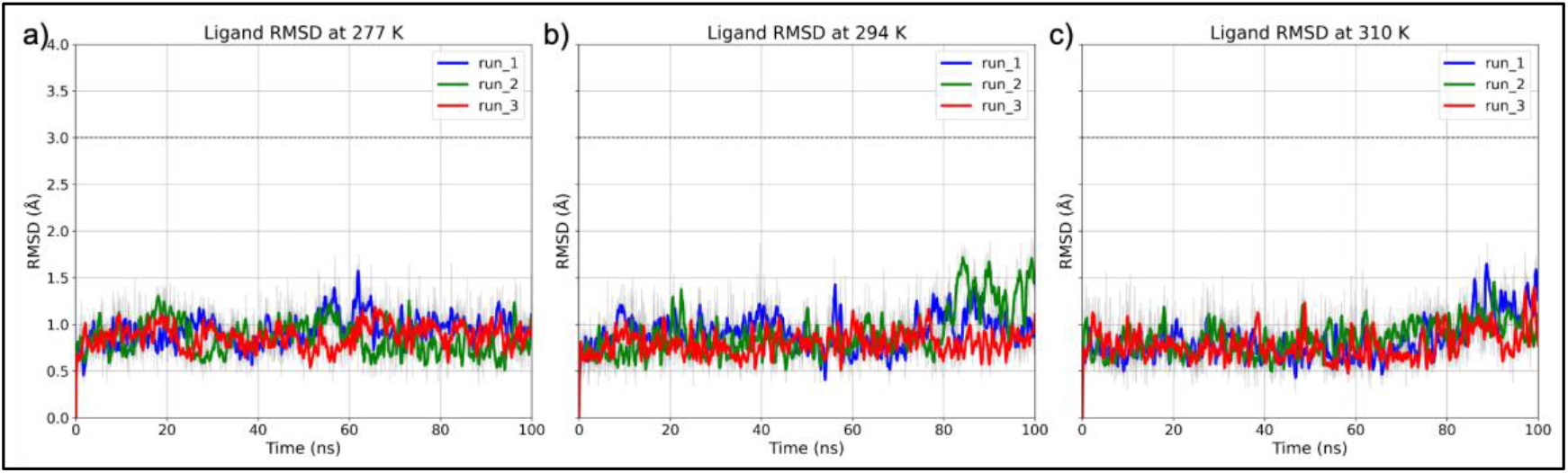
Temporal evolution of ligand RMSD during molecular dynamics simulations. Root-mean-square deviation (RMSD) trajectories of UCB-F within the SV2C complex are shown for simulations conducted at 277 K (a), 294 K (b), and 310 K (c). For each temperature, three independent simulation replicates (run_1, run_2, and run_3) are displayed as blue, green, and red lines, respectively. RMSD values were calculated over 100 ns trajectories with respect to the initial binding pose.

While RMSD analysis provides information on the overall structural stability of the complex, it does not capture local interactions that are important for maintaining ligand binding. In this system, hydrogen bonds between UCB-F and SV2C have been identified as key stabilizing interactions based on the predicted binding mode. Therefore, we performed additional analysis of the simulation trajectories to assess the behaviour of these hydrogen bonds under different temperature conditions. One of the hydrogen bonds remained relatively stable across all temperatures, whereas the other (Figure S) exhibited temperature-dependent fluctuations (Figure 3). As shown in Figure 3, the lifetime of this hydrogen bond varied with temperature. At 277 K, the hydrogen bond persisted for longer periods, with a mean lifetime of 11.86 ns. As the temperature increased to 294 K and 310 K, the lifetimes decreased, with the shortest mean lifetime of 1.03 ns observed at 310 K.

**Figure 3.**
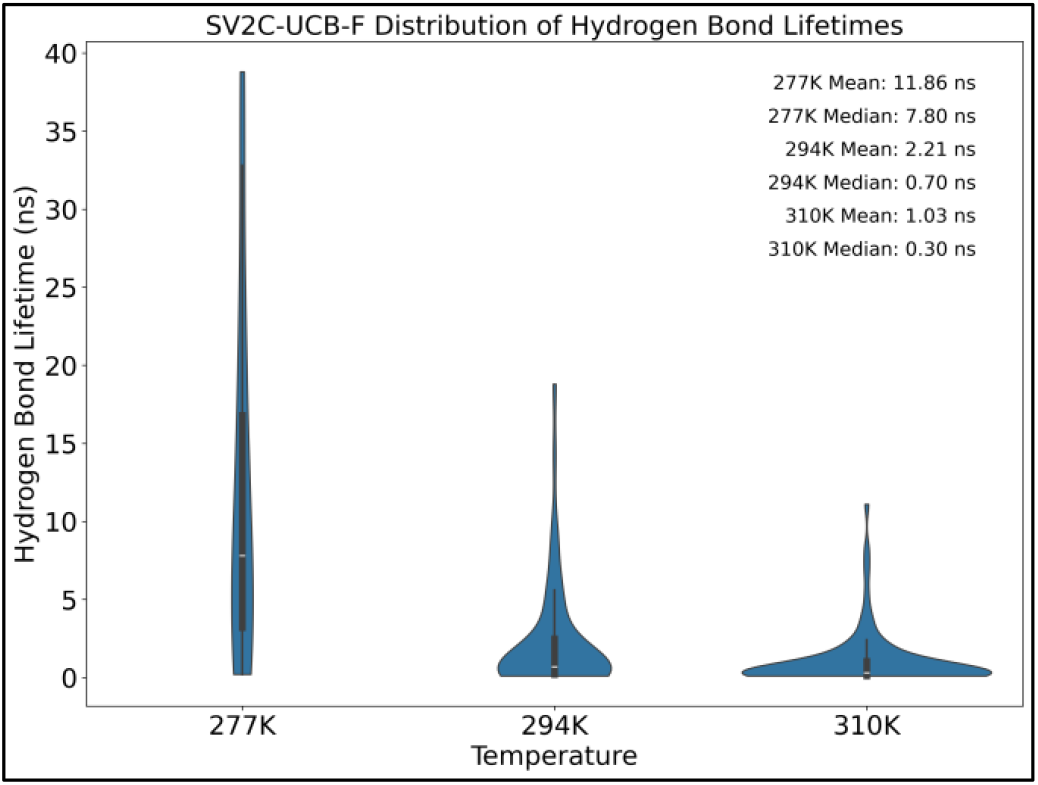
Violin plots depicting the distribution of hydrogen bond lifetimes between the UCB-F ligand and SV2C. Data are shown for three different temperatures: 277 K, 294 K, and 310 K. The distributions illustrate the stability and persistence of hydrogen bonding interactions under varying thermal conditions.

This reduction in hydrogen bond lifetime at elevated temperatures reflects a weakening of specific stabilizing interactions within the binding site, which may contribute to lower binding affinity of UCB-F to SV2C under these conditions. Experimental data indicate that UCB-F exhibits reduced binding affinity at higher temperatures. Therefore, our simulation results provide a structural perspective that helps to explain the experimentally observed temperature dependence of ligand binding, suggesting that decreased hydrogen bond stability is associated with weakened binding interactions.

### Radiolabelling of [^18^F]UCB-F

The radiolabeling of [^18^F]UCB-F was successfully carried out through a nucleophilic substitution process, where the respective mesylate precursor was reacted with [^18^F]fluoride in a one-pot synthesis. This process utilized kryptofix K_2.2.2_ and potassium carbonate (K_2_CO_3_) to facilitate the reaction, as shown in Scheme 1. Several solvents, including acetonitrile, dimethylformamide (DMF) and dimethyl sulfoxide (DMSO), were evaluated at varying reaction temperatures to optimize the yield. The combination of acetonitrile as the solvent and a reaction temperature of 100°C maintained for 10 minutes proved to be the most effective, resulting in the highest radiochemical yield for the synthesis.

**Scheme 1.**
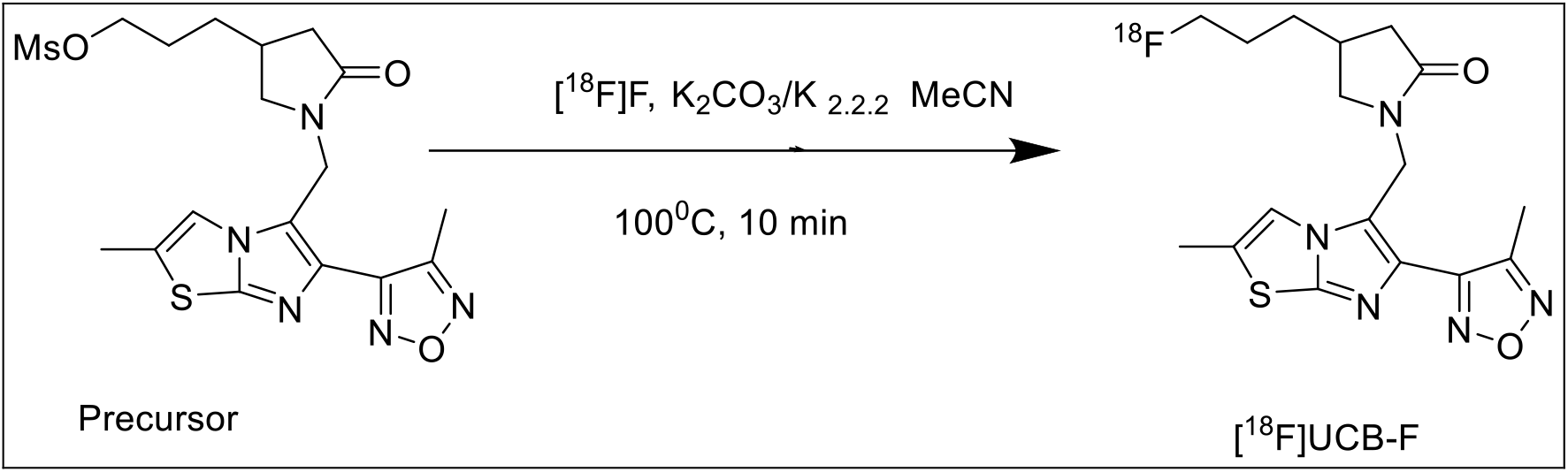

Following synthesis, the radiolabeled [^18^F]UCB-F was purified using high-performance liquid chromatography (HPLC). From approximately 10 minutes of cyclotron irradiation at 35 μA, we obtained over 3.0 GBq of the final product. The overall radiochemical yield, non-decay corrected, was consistently above 35%. [^18^F]UCB-F achieved radiochemical purities exceeding 99%, demonstrating the high purity of the preparations. The entire radiosynthesis workflow including [^18^F]-fluorination, HPLC purification, solid-phase extraction (SPE) isolation, and formulation— was completed within 75 minutes.

### Autoradiography studies in rat and non-human primate brain tissue

[^18^F]UCB-F binding was evaluated by autoradiography in brain sections of *n*=3 rats and n=2 NHP (Figure 4). The distribution of [^18^F]UCB-F binding was comparable between corresponding brain regions in the rat and NHP and follows the expression profile of SV2C in the brain ^12-14^. The highest binding levels were observed in the brainstem, deep cerebellar nuclei (DCN) and basal ganglia, particularly within the striatum, globus pallidus and substantia nigra (Table 1). The lowest binding was observed in cerebral and cerebellar cortex as well as the hippocampus, where immunostaining experiments have shown to express low levels of SV2C. With co-incubation of 10 µM cold UCB-F, only <10% of the [^18^F]UCB-F binding was non-displaceable in the rat brain regions expressing high SV2C (basal ganglia, brainstem, DCN), and ~20% in the cerebral and cerebellar cortex. In the NHP, the level of [^18^F]UCB-F binding that was not displaced by 10 µM cold UCB-F was ~20% in the basal ganglia and ~40% in the cortical regions. We believe this higher level of non-displaceable binding in the NHP sections to be due to the thickness of the tissue, since the NHP sections were 20µm thick, whereas the rat sections were 12µm.

**Table 1.** Specific binding (Bq/mm2) of [^18^F]UCB-F to synaptic vesicle glycoprotein 2C (SV2C), measured in rat and non-human primate brains.

|  | Rat |  |  | Non-Human Primate |  |  |
| --- | --- | --- | --- | --- | --- | --- |
| Brain region | n | Specific binding<br>Bq/mm<sup>2</sup> | ±SEM | n | Specific binding<br>Bq/mm<sup>2</sup> | ±SEM |
| Cortex | 3 | 0.7 | 0.1 | 2 | 2.6 | 0.2 |
| Hippocampus | 3 | 0.9 | 0.1 |  |  |  |
| Striatum | 3 | 3.6 | 0.3 |  |  |  |
| Putamen |  |  |  | 2 | 4.5 | 0.6 |
| Caudate nucleus |  |  |  | 2 | 4.2 | 1.1 |
| Ventral pallidum | 3 | 5.2 | 0.3 |  |  |  |
| Globus pallidus |  |  |  | 2 | 5.9 | 0.6 |
| Substantia nigra | 2 | 5.1 | 1.5 | 1 | 5.4 | 0.0 |
| Deep cerebellar nuclei | 3 | 2.8 | 0.3 |  |  |  |
| Cerebellum | 3 | 0.9 | 0.1 | 2 | 0.9 | 0.1 |
| Brainstem | 3 | 4.1 | 0.1 |  |  |  |

Specific binding is defined as the difference between the total binding (0.15 MBq/mL [<sup>18</sup>F]UCB-F) and the binding remaining in the presence of an excess amount (10 µM) of unlabelled UCB-F. Values are presented as mean specific binding (Bq/mm<sup>2</sup>) ±standard error of the mean (SEM) from the indicated number (n) of individuals. Quantification of the rat striatum consisted of the average of the putamen and caudate nucleus.

**Figure 4.**
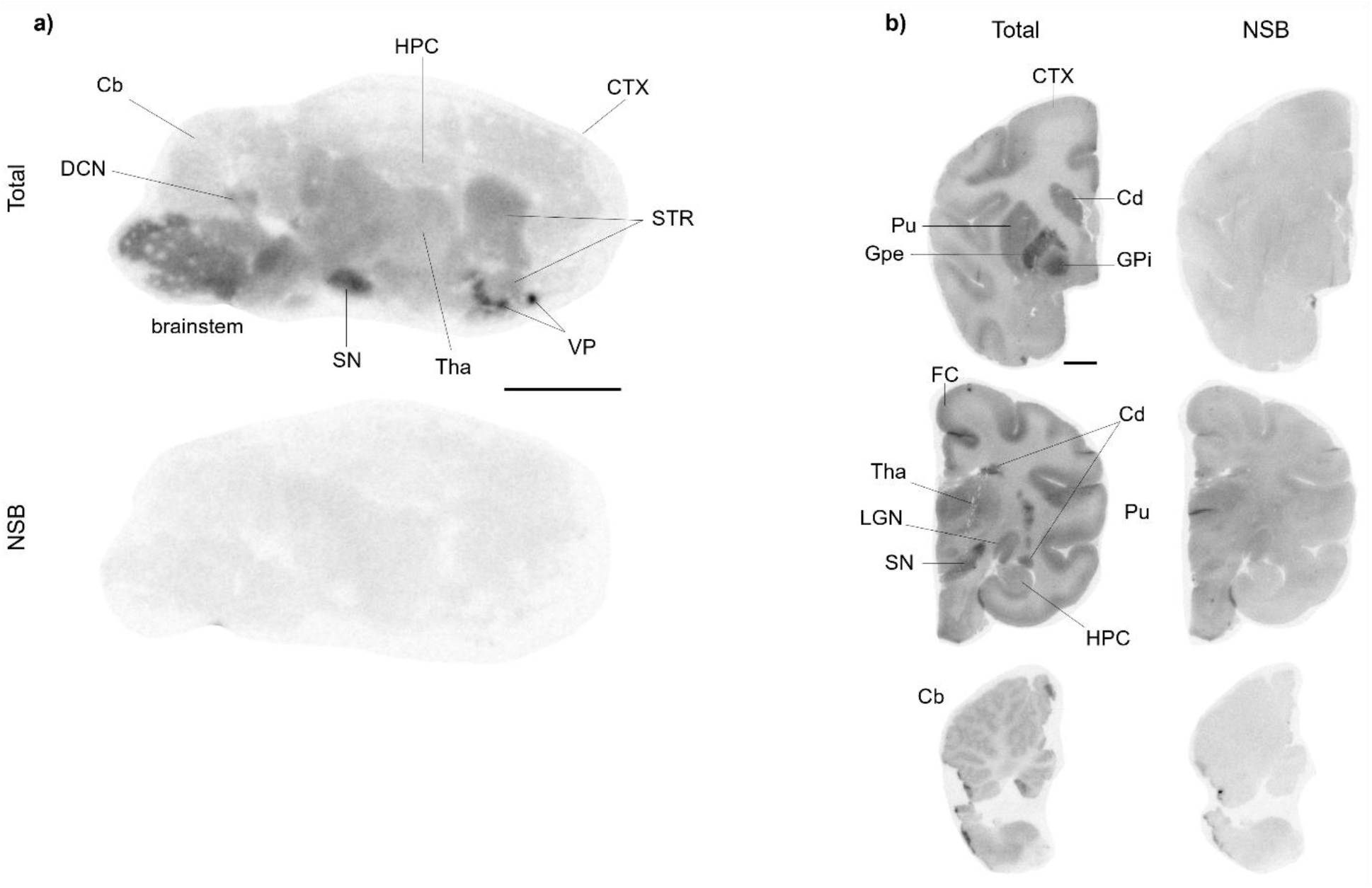
Representative autoradiograms of [^18^F]UCB-F binding to synaptic vesicle glycoprotein 2C (SV2C). The images represent (a) rat brain sagittal sections (0.15 MBq/mL [^18^F]UCB-F, 25.8 GBq/µmol) and (b) non-human primate coronal sections (0.15 MBq/mL [^18^F]UCB-F, 15.9 GBq/µmol), alone (Total) and co-incubated with excess amount (10 µM) of unlabelled UCB-F (non-specific binding, NSB). CTX, cortex; STR, striatum; Cd, caudate nucleus; Pu, putamen; VP, ventral pallidum; GPi, internal globus pallidus; GPe, external globus pallidus; Tha, thalamus; LGN, lateral geniculate nucleus; HPC, hippocampus; SB, substantia nigra; Cb, cerebellum; DCN, deep cerebellar nuclei. Line scales: 5 mm.

### In vitro competition studies

In vitro competition binding (n=3) demonstrates a clear temperature dependence for UCB-F, with SV2C affinity decreasing by about an order of magnitude from 4°C to 37°C, consistent with shorter hydrogen-bond lifetimes in the binding pocket at higher temperature. At 4°C, human UCB-F shows high affinity (pKi 8.4 with [^3^H]UCB101275-1), whereas at 37°C SV2C affinity falls to pKi ~6.8–7.3 across species and radioligands (Table 2). In contrast, at 37°C UCB-F exhibits minimal binding to human SV2A and SV2B (pKi <6 with [^3^H]padsevonil), establishing a robust SV2C-selective profile under physiologically relevant conditions (Table 2). Together, these data indicate that cooling artificially inflates apparent potency (pKi ~8.3–8.4 at 4°C), while at 37°C UCB-F maintains selective SV2C engagement in the low-to mid–double-digit nanomolar range, with at least a 20–200× selectivity window over SV2A/SV2B depending on species and radioligand; the 4°C SV2C value is reported elsewhere^15^.

**Table 2.** In vitro pKi (n=3) of UCB-F to SV2 proteins measured at 4°C and 37°C.

| Radioligand | hSV2C<br>(4°C) | hSV2C<br>(37°C) | rSV2C<br>(37°C) | pSV2C<br>(37°C) | hSV2A<br>(37°C) | hSV2B<br>(37°C) |
| --- | --- | --- | --- | --- | --- | --- |
| [<sup>3</sup>H]UCB101275-1<sup>15</sup> | 8.4 | 7.3 | 6.9 | 7.0 |  |  |
| [<sup>3</sup>H]UCB-1A | - | 7.3 | 6.7 | 6.8 |  |  |
| [<sup>3</sup>H]padsevonil | - | 6.8 |  |  | <6 | <6 |

[<sup>3</sup>H]UCB-1A = [<sup>3</sup>H]UCB4297

### PET studies with [^18^F]UCB-F in non-human primates

Two cynomolgus monkeys, one female (NHP1) and one male (NHP2), were examined with [^18^F]UCB-F, as detailed in Table 3. The injected radioactivity of [^18^F]UCB-F was measured 160 MBq and 144 MBq, with an injected mass of 2.3 and 3.4 µg. Summated PET images for the two baseline studies are presented in Figure 5. Under baseline condition, [^18^F]UCB-F showed whole-brain uptake peaking at 3.4 SUV for NHP1 and 3.0 SUV for NHP2 (Figures 6). An initial rapid increase in radioligand uptake was observed across the entire brain, with minimal SUV variation among regions (Figure 7). The radioligand exhibited rapid washout across all regions, indicating reversible kinetics.

**Table 3.**
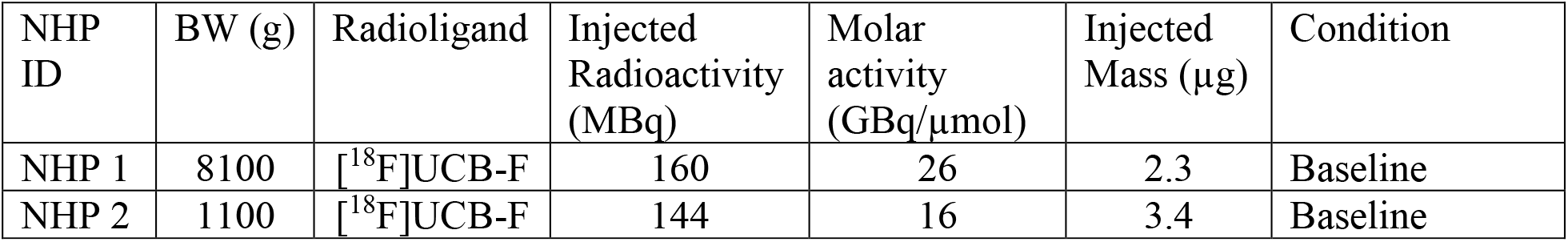

**Figure 5.**
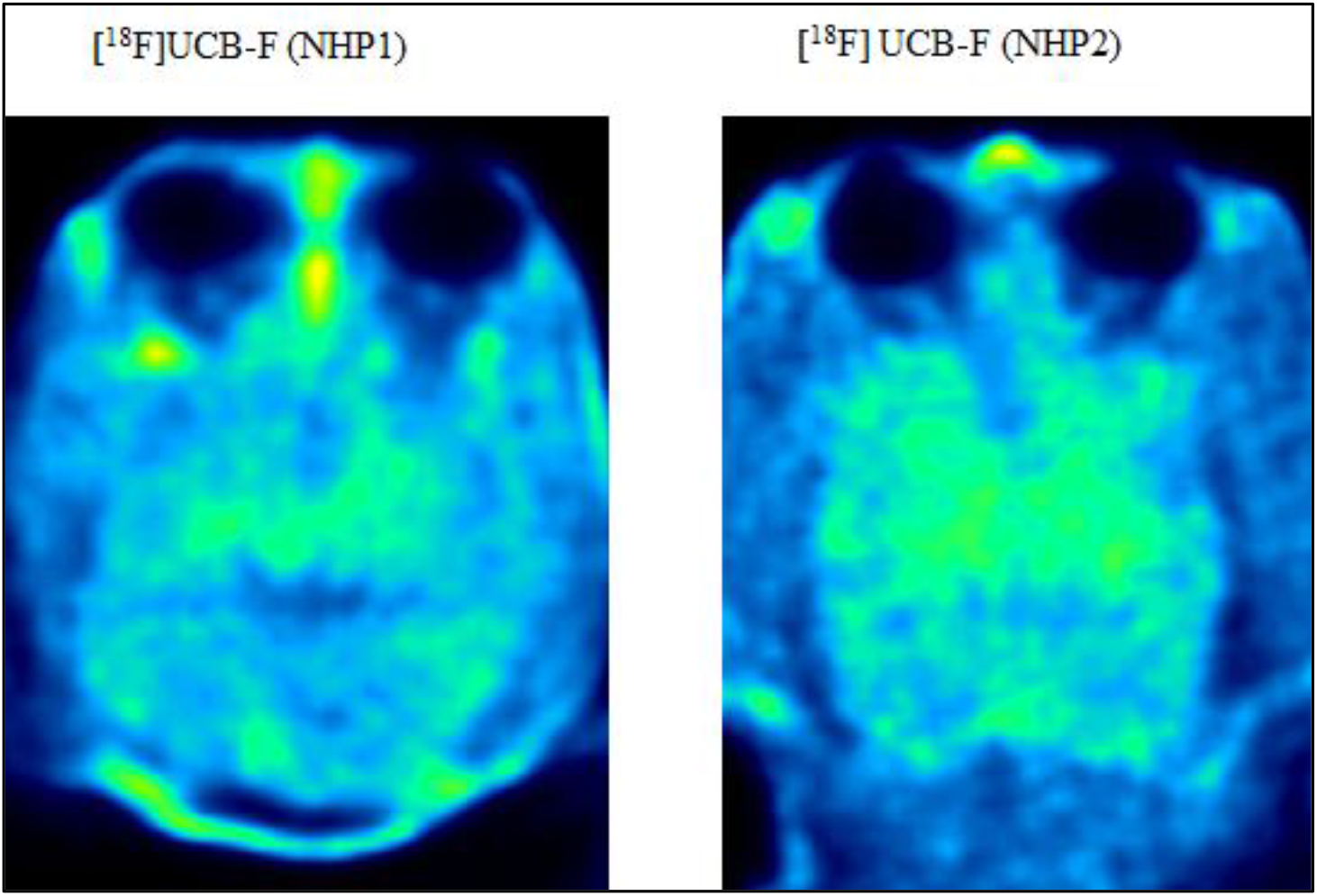
PET images in NHP1 and NHP 2 after administration of [^18^F]UCB-F.

**Figure 6.**
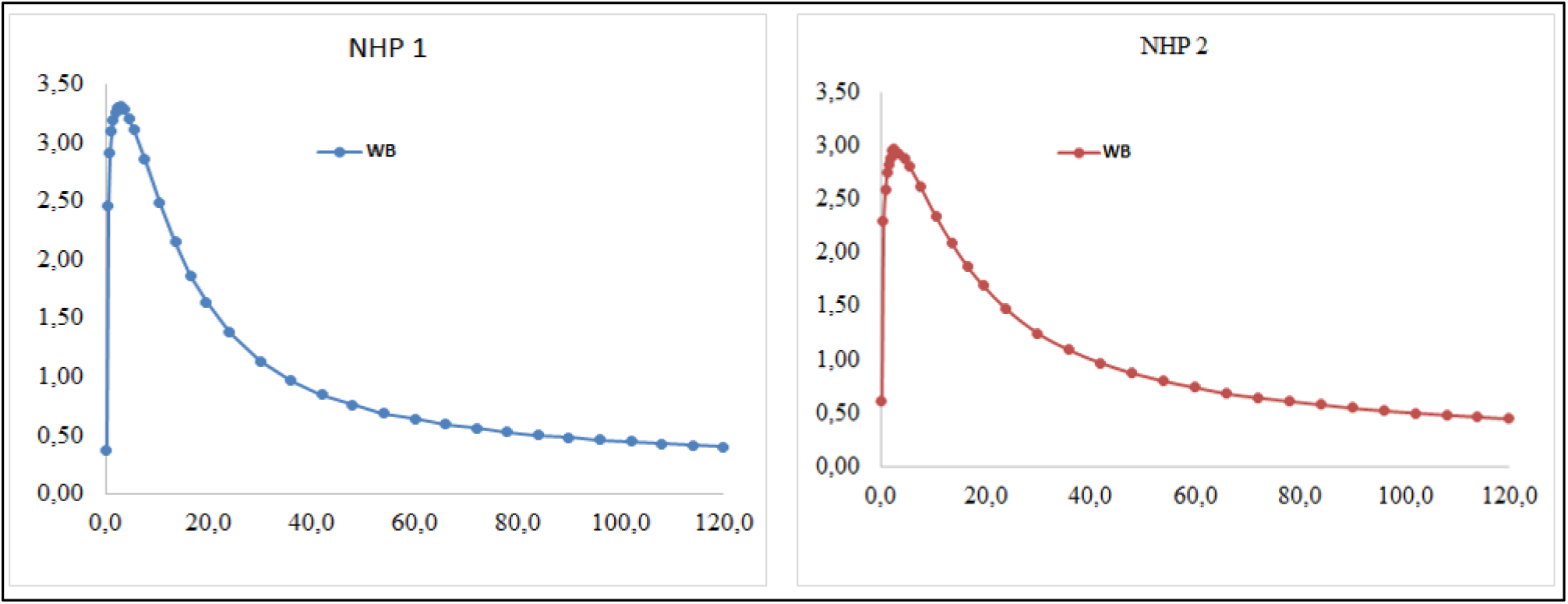
Time activity curves representing whole brain uptake of [^18^F]UCB-F in NHP 1 and in NHP 2 at baseline condition.

**Figure 7.**
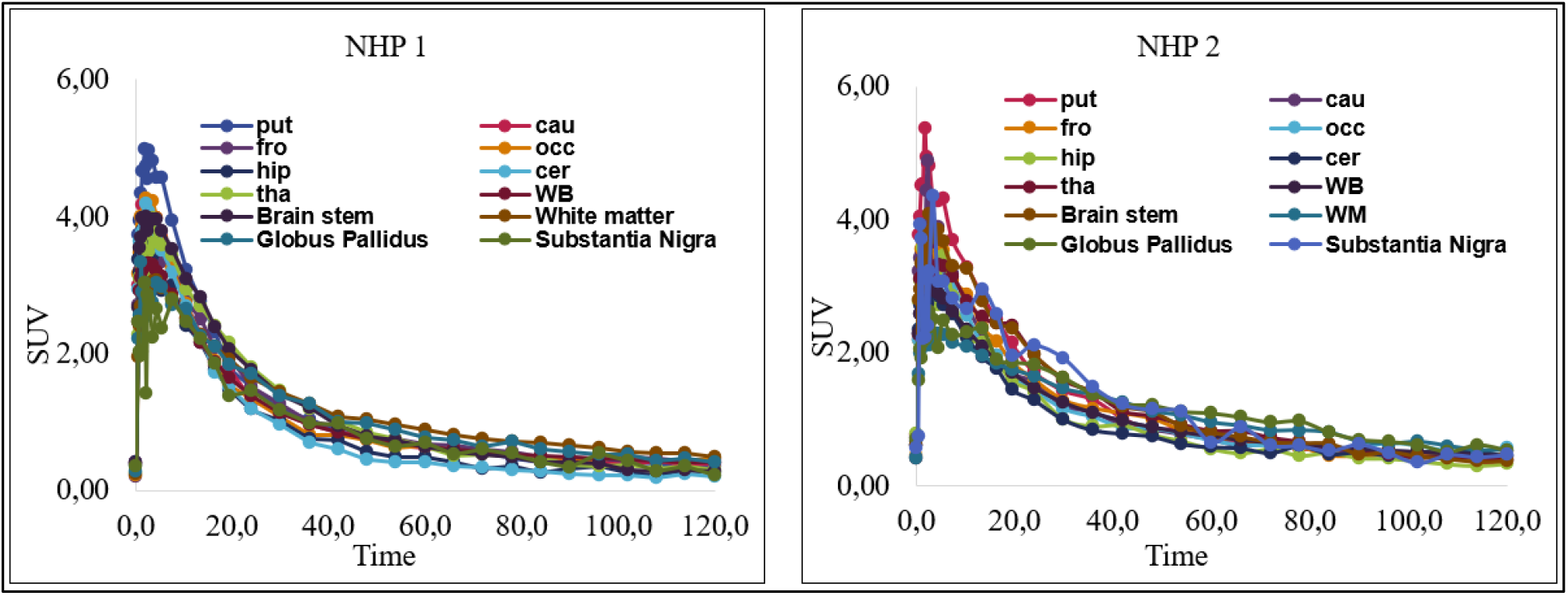
Time activity curves representing regional brain uptake of [^18^F]UCB-F in NHP 1 and in NHP 2 at baseline condition.

### Radiometabolite analysis

After deproteinization, over 95% of the plasma radioactivity was effectively recovered in acetonitrile. HPLC analysis following the injection of [^18^F]UCB-F into plasma showed the compound eluting at a retention time of 6.5 minutes. Initially, the parent compound was most abundant at 2 minutes, accounting for approximately 92%, but this decreased to around 2% at 45 minutes during PET studies, as reflected in the percentage of unchanged radioligand across both PET experiments (Figure 8). Additionally, several more polar radiometabolite peaks appeared before the parent peak during elution. The identity of [^18^F]UCB-F was confirmed through coinjection with its non-radioactive counterpart.

**Figure 8.**
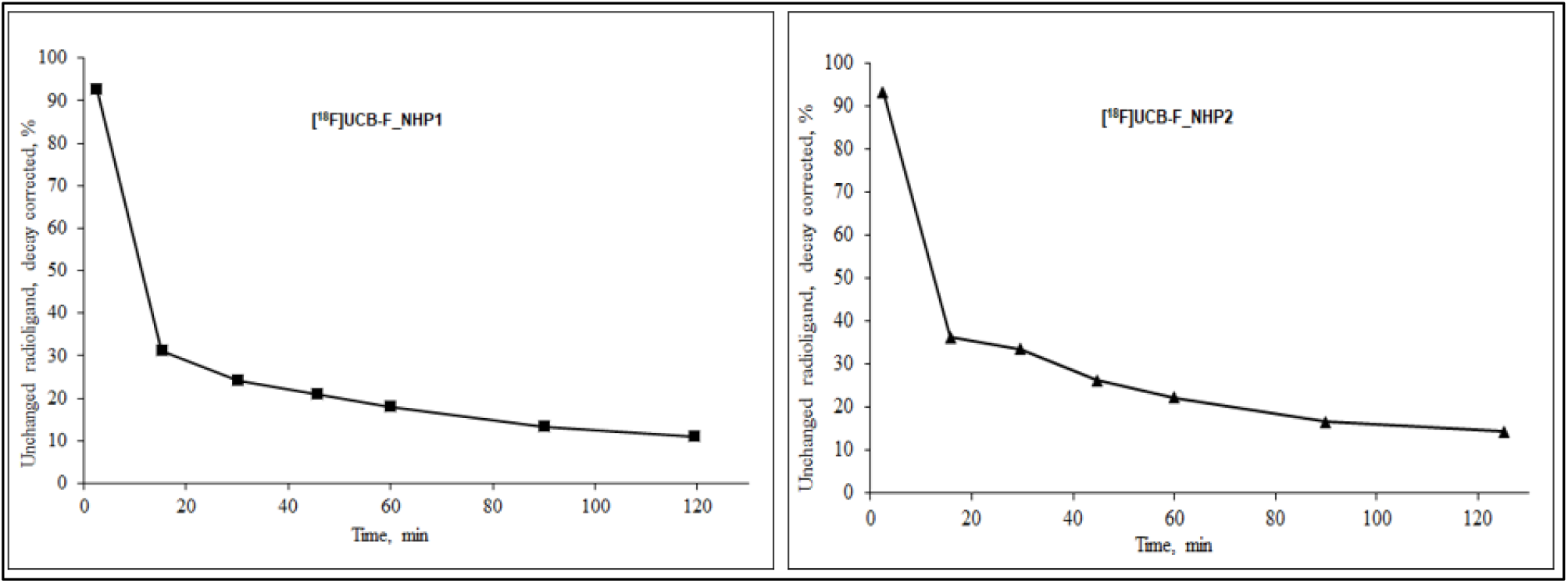
The *in vivo* metabolism of [^18^F]UCB-F is shown as the relative plasma composition at baseline condition in NHP1 and NHP2

## Conclusion

[^18^F]UCB-F was successfully radiolabelled with high incorporation yield. *In vitro*, it showed specific binding on tissue slices from rodent and NHP brains. However, *in vivo*, [^18^F]UCB-F displayed rapid washout from the brain, lack of regional brain distribution and rapid metabolism. The discrepancy between autoradiography and *in vivo* PET studies is likely explained by the decrease in the affinity of [^18^F]UCB-F to SV2C that was observed at 37°C compared with the affinity measured at 4°C. *In silico* modelling studies suggested that the effect of temperature on the affinity of UCB-F to SV2C can be explained by the decreased hydrogen bond stability of the binding pose. Overall, these findings indicate that [^18^F]UCB-F is not a suitable PET radioligand for imaging SV2C and additional work is ongoing to identify a potential candidate.

## Materials and Methods

### *In silico* calculations including machine learning

The initial structure of SV2A was obtained from the Protein Data Bank using PDB ID: 8UO9^16^ and prepared using the Protein Preparation Wizard in Schrödinger. The UCB-F ligand was positioned in the binding site by aligning it with the co-crystallized reference ligand, UCB-2500. The OPM (Orientations of Proteins in Membranes) webserver^17^ was used to determine the appropriate orientation of the protein within a lipid bilayer.

The SV2C–UCB-F complex was generated via homology modelling using the SV2A–UCB-F structure as a template. System construction for SV2C was carried out using Schrödinger’s System Builder module. The complex was embedded in a POPC lipid bilayer, solvated with TIP3P water in an orthorhombic box with a 10 Å buffer between the protein and the box edges, and neutralized with Na^+^ and Cl^−^ ions to achieve a physiological salt concentration of 0.15 M. A 10 ns equilibration simulation was performed to relax the system prior to production runs.

Production molecular dynamics simulations were conducted at three temperatures (277 K, 294 K, and 310 K) to investigate temperature-dependent effects on binding dynamics. All simulations were performed using the OPLS4 force field implemented in Desmond (Schrödinger 2023-2)^18^. Production runs were extended to 100 ns to ensure adequate conformational sampling. For each temperature condition, three independent replicates were performed with different initial random seeds. Trajectory analyses were conducted using Desmond’s built-in tools, complemented by custom Python scripts.

### Radiochemistry

#### General

The non-radioactive reference standard UCB-F (4-(3-(fluoropropyl)-1-((2-methyl-6-(4-methyl-1,2,5-oxadiazol-3-yl)imidazo[2,1-b]thiazol-5-yl)methyl)pyrrolidin-2-one), as well as the corresponding precursor 3-(1-((2-methyl-6-(4-methyl-1,2,5-oxadiazol-3-yl)imidazo[2,1-b]thiazol-5-yl)methyl)-5-oxopyrrolidin-3-yl)propyl methanesulfonate—were synthesised by Advinus Therapeutics, Bangalore-560058, India. All other chemicals and reagents were sourced from commercial suppliers and used without further purification. Solid-phase extraction (SPE) cartridges, specifically SepPak QMA Light and SepPak C18 Plus, were purchased from Waters, Milford, Massachusetts, USA. Before use, the C18 Plus cartridges were activated by rinsing with 10 mL of ethanol followed by 10 mL of sterile water. The SepPak QMA Light cartridges were activated using 10 mL of 0.5 M K2CO3 solution, followed by rinsing with 15 mL of 18 MΩ water. Fluorine-18 fluoride ([18F]F) was produced at Karolinska Hospital in Stockholm, Sweden, using a 16.4 MeV GEMS PET trace cyclotron (GE, Uppsala, Sweden). Radiolabeling was performed using a custom-made semi-automated or automated synthesis module. Liquid chromatography (LC) analysis was carried out on a Merck-Hitachi gradient pump with a Merck-Hitachi L-4000 variable wavelength UV detector. LC-MS analyses were performed using either a Waters Quattro Premier micro mass spectrometer or a Waters SQD 3001 single quadrupole mass spectrometer, both connected to Waters Acquity UPLC systems.

### Production of [^18^F]Fluoride ([^18^F]F^-^)

The production of ^18^F-/K_2_CO_3_/Kryptofix_2.2.2_ complex, an organic cation/inorganic anion salt was carried out following a previously published method^19^. To generate [18F]F−, a GEMS PETtrace cyclotron was used to irradiate 18O-enriched water with 16.4 MeV protons via the 18O(p,n)18F reaction. The produced [18F]F− was separated from the 18O-H2O using a conditioned SepPak QMA Light anion exchange cartridge, which was then washed with a solution containing potassium carbonate (13 µmol, 1.8 mg) and kryptofix 2.2.2 (26 µmol, 9.8 mg) in 85 µL of 18 MΩ water and 2 mL of acetonitrile (MeCN). This mixture was transferred into a reaction vessel (either 10 mL or 4 mL). The solvents were evaporated at 140°C under a continuous flow of nitrogen/helium (70 mL/min) for 10 minutes, resulting in a dry complex of [18F]F−/K2CO3/K2.2.2. After evaporation, the residue was cooled to room temperature.

### Synthesis of [^18^F]UCB-F (4-(3-[^18^F]fluoropropyl)-1-((2-methyl-6-(4-methyl-1,2,5-oxadiazol-3-yl)imidazo[2,1-b]thiazol-5-yl)methyl)pyrrolidin-2-one)

To the dry complex of ^18^F-F-/K_2_CO_3_/K_2.2.2_., corresponding mesylate-precursor (1 mg, 0.005 mmol) in acetonitrile (500 µL) was added at 100°C and left for 10 min to produce <u>[^18^F]UCB-F</u>. The reaction mixture was cooled to room temperature (RT) and diluted with water (2.5 mL) to a total volume of 3 mL before being injected into a semi-preparative reverse-phase XBridge HPLC column (C18, 10 mm × 250 mm, 5 µm) for purification. The column outlet was connected to a UV absorbance detector (λ = 254 nm) in series with a GM-tube for radioactivity detection. Elution was carried out with a mobile phase consisting of CH_3_CN and ammonium formate (0.1M) in a ratio of 24:76 at a flow rate of 6 mL/min. This process yielded a radioactive fraction corresponding to pure [^18^F]1/[^18^F]2, with a retention time (t_R_) of 20-22 minutes and collected in a bottle containing sterile water (50 mL). The resulting mixture was further purified by passing through a preconditioned SepPak SPE (Oasis HLB 3cc) cartridge followed by washing with sterile water (10 mL). Isolated [^18^F]1/[^18^F]2 was eluted with 1 mL of ethanol into a sterile vial containing 9 mL of sterile saline. The formulated product was then filtered through a Millipore Millex® GV sterile filter unit (0.22 μm) for further application in PET.

#### *In vitro* Autoradiography

Sectioning of the rat and NHP tissue and autoradiography assays were performed, as previously described^20^ at the Autoradiography Core Facility, Department of Clinical Neuroscience, Division of Imaging Core Facilities and Center for Imaging Research (CIR), at the Karolinska Institutet. Studies using NHP brain tissue were approved by the Animal Ethics Committee of the Swedish Animal Welfare Agency (registration no. 4820/06-600 and 399/08). Sections from the brains of 2 adult female cynomolgus monkeys (Macaca Fascicularis) were used in this study. Coronal sections from the snap-fozen brains of *n*=3 adult naïve Sprague Dawley rats were sectioned at 12 μm thickness, and NHP Brains were cut at 20 µm using a cryomicrotome (CM1860, Leica Biosystems, Nußloch, Germany), thaw-mounted onto glass slides and stored at −20 °C until the autoradiography assay was performed. For the [^18^F]UCB-F binding autoradiography assay, all sections were first pre-incubated for 20 minutes at room temperature with binding buffer (50 mM Tris HCl, pH 7.4, containing 120mM NaCl, 5mM KCl, 2mM CaCl2, 1mM MgCl2), then incubated at room temperature with [^18^F]UCB-F (0.15 MBq/mL) for 55 minutes. The rat and NHP autoradiography assays were performed in separate days. The molar activity measured was 25.8 GBq/µmol in the rat autoradiography assay and 15.9 GBq/µmol in the NHP autoradiography assay. Non-specific binding was determined by co-incubation with 10 μM of unlabelled UCB-F. After incubation, sections were washed 3 x 5 min at 4 °C, in 50 mM Tris HCl, pH 7.4, briefly dipped in ice-cold distilled water, then dried in a heat plate before being exposed to storage phosphor screens (BAS-IP SR2025, Fujifilm, Tokyo, Japan) overnight.

Radioactivity was detected and quantified with an Amersham Typhoon FLA-9500 phosphor imaging scanner (Cytiva, Marlborough, MA, USA). Autoradiograms were analyzed using Multi Gauge 3.2 phosphor imager software (Fujifilm, Tokyo, Japan). The measured photo-stimulated luminescence (PSL)/mm^2^ values were converted into decay-corrected radioactivity units based on the signal intensity from standard quantities (1.5 kBq to 2.9 kBq), diluted from the respective [^18^F]UCB-F batch, and pipetted (20 µL per standard) onto filter papers exposed in each storage phosphor plate. Binding values from two replicates of total and non-specific conditions from each individual were plotted using GraphPad Prism v10 (GraphPad Software, Boston, MA, USA) and specific binding was determined by subtracting the binding signal in the presence of 10 µM unlabelled (cold) UCB-F from the total binding.

### Study Design in non-human primates (NHP), PET procedure and quantification

This research study adhered to ethical standards and received approval from the Animal Ethics Committee in Stockholm, administered by the Swedish Board of Agriculture (10367-2019). The study also complied with the guidelines outlined in *”Guidelines for Planning, Conducting, and Documenting Experimental Research”* (Dnr 4820/06-600) of Karolinska Institutet. The non-human primates (NHPs) used in this study were housed at the Astrid Fagraeus Laboratory, Comparative Medicine, in Solna, Sweden. Two male cynomolgus NHPs were selected for participation. The study involved performing brain PET scans on both NHPs under baseline conditions.

At the Astrid Fagraeus Laboratory, each non-human primate (NHP) received ketamine hydrochloride at 10 mg kg-1 by intramuscular injection to induce anesthesia. An inflatable cuff cuff was inflated on the delivery hose until capnography confirmed clear alveolar waves. Endotracheal intubationmaintained anesthesia with a blend of sevoflurane, oxygen and medical air. A three-point head holder fixed the skull, while an esophageal probe fed a Bair Hugger 505 unit (Arizant Healthcare, MN) to provide warmth. Heart rate, blood pressure, respiratory rate and SpO2 were recorded on a bedside monitor, and saline dripped continuously to correct volume losses.

Positron emission tomography was performed on a LFER PET-CT system (Mediso)^21^. [^11^C]UCB-1A was administered as bolus, followed by the administration of 2 mL NaCl. PET data were acquired in list mode for 123 min and reconstructed with a series of frames of increasing duration (9 x 20 sec, 3 x 60 sec, 5 x 180 sec, 17 x 360 sec). The reconstruction was performed using the Tera-Tomo 3D algorithm (with 10 iterations and 9 subsets), including detector modelling with attenuation and scatter correction based on a 3-component material map. Volumes of interest (VOIs) were delineated on an MR image of the animal, which was manually coregistered with the average PET image using the FUSION tool in PMOD software (version 4.2; Bruker). Selected VOIs were caudate, putamen, globus pallidus, substantia nigra, and cerebellum. Radioactivity concentration was expressed as SUV. Venous blood samples were collected at 2, 15, 30, 45, 60, 90, and 120 min for radioactivity measurements and radiometabolite analysis.

Magnetic resonance imaging was performed on a General Electric DISCOVERY MR750 3.0 T scanner (Milwaukee WI, USA). A T1-weighted sequence was acquired to facilitate co-registration with PET and delineate cortical and subcortical regions. Data analysis began with the investigator hand-drawing regions of interest (ROIs) on the MRIs of each non-human primate, outlining the entire brain as well as the cerebellum, caudate, putamen, thalamus, frontal cortex, occipital cortex, and hippocampus. Next, the PET scans covering the full imaging session were precisely aligned to each animals structural MRI. The alignment parameters were then transferred to the dynamic PET dataset, allowing time-activity curves to be extracted from all specified brain regions. Finally, the average standardized uptake value (SUV) was computed for every delineated ROI.

### Radiometabolite analysis

Radiometabolites present in the blood plasma of non-human primates (NHP) were analyzed using a previously published method^22^. Throughout the PET imaging session, the proportions of radioactivity attributable to [^18^F]UCB-F and its radioactive metabolites in NHP plasma were quantified using a reverse-phase high-performance liquid chromatography (HPLC) system. This system employed a UV detector set at 254 nm in conjunction with a radioactive detector. Arterial blood samples, ranging from 1.0 to 3.0 mL, were manually collected at various time points, including 2.5, 15, 30, 45, 60, 90, and 120 minutes after injection of [^18^F]UCB-F. Post-collection, blood samples were centrifuged at 4000 rpm for 2 minutes to isolate plasma, which was then diluted 1.4-fold with acetonitrile and centrifuged at 6000 rpm for 4 minutes. The supernatant was separated from the pellet and further diluted with 3 mL of water. For the separation of radiometabolites from the unchanged [^18^F]UCB-F, a high-performance liquid chromatography (HPLC) system was employed. The setup included an Agilent 1200 series binary pump connected to a Rheodyne 7725i manual injection valve with a 5.0 mL loop, and a radiation detector (Oyokoken S-2493Z) enclosed within a 50 mm lead shield. A semi-preparative ACE 5µm C18 HL column (250 × 10 mm) was used for chromatographic separation, utilizing gradient elution. The mobile phase consisted of acetonitrile (A) and 0.1 M ammonium formate (B), flowing at 5.0 mL/min. The gradient program involved: 0-7.0 minutes, changing from 40:60 to 90:10 (A/B) v/v; 7.0-9.0 minutes, maintaining at 90:10; and 9.0−9.1 minutes, returning from 90:10 to 40:60 (A/B) v/v. The radioactive peaks were integrated, with the peak corresponding to [^18^F]UCB-F expressed as a percentage of the total detected radioactivity. To assess the system’s recovery rate, a 2 mL aliquot of the collected eluate was measured and divided by the total radioactivity.

## Author contributions

The manuscript was written through contributions of all authors. All authors have given approval to the final version of the manuscript.

## Institutional Review Board Statement

The study was approved by the Animal Ethics Committee of the Swedish Animal Welfare Agency and was performed according to “Guidelines for planning, conducting and documenting experimental research” (Dnr 4820/06-600)

## Data Availability Statement

All the supporting data are stored at Karolinska Institutet archive.

## Acknowledgments

The authors would like to thank the members of the Core Facilities of Radiopharmacy, PET (Brain molecular imaging centre) and autoradiography for their excellent technical assistance in the PET and ARG experiments.

